# Structural basis of recruitment of RBM39 to DCAF15 by a sulfonamide molecular glue E7820

**DOI:** 10.1101/758409

**Authors:** Xinlin Du, Oleg Volkov, Robert Czerwinski, HuiLing Tan, Carlos Huerta, Emily Morton, Jim Rizzi, Paul Wehn, Rui Xu, Deepak Nijhawan, Eli Wallace

## Abstract

E7820 and indisulam are two examples of aryl sulfonamides that recruit RBM39 to Rbx-Cul4-DDA1-DDB1-DCAF15 E3 ligase complex, leading to its ubiquitination and degradation by the proteasome. In order to understand their mechanism of action, we carried out kinetic analysis on the recruitment of RBM39 to DCAF15 and solved a crystal structure of DDA1-DDB1-DCAF15 in complex with E7820 and the RRM2 domain of RBM39. E7820 packs in a shallow pocket on the surface of DCAF15 and the resulting modified interface binds RBM39 through the α1 helix of RRM2 domain. Our kinetic studies revealed that aryl sulfonamide and RBM39 bind to DCAF15 in a synergistic manner. The structural and kinetic studies confirm aryl sulfonamides as molecular glues in the recruitment of RBM39 and provide a framework for future efforts to utilize DCAF15 to degrade other protein of interests.

## Introduction

E7820 and indisulam (Fig. 1a) are aryl sulfonamide agents that have anti-proliferative effects on certain cell lines *in vitro* and anti-tumor effects on tumor xenografts in mice (Owa et al., 1999; Semba et al., 2004). Although the molecular mechanism underpinning these activities was unknown until recently, both E7820 and indisulam have been tested extensively in clinical trials targeting a variety of cancer indications either as a single agent or in combination with other treatments (Siegel-Lakhai et al. 2008; Mita et al., 2011; Milojkovic Kerklaan et al. 2016). Recently two studies reported that indisulam and other aryl sulfonamides recruit RNA-splicing factor RBM39 to DCAF15, a substrate adaptor in the Rbx-Cul4-DDB1-DCAF15 E3 ligase (Han et al., 2017; Uehara et al. 2017). Proteasomal degradation of RBM39 leads to aberrant processing of pre-mRNA in hundreds of genes, primarily reflected by intron retention and exon skipping. (Han et al., 2017). Hence, indisulam and other aryl sulfonamides such as E7820, tasisulam, and chloroquinoxaline were collectively referred to as SPLicing inhibitor sulfonAMides, or SPLAMs (Han et al., 2017).

**Figure 1.**
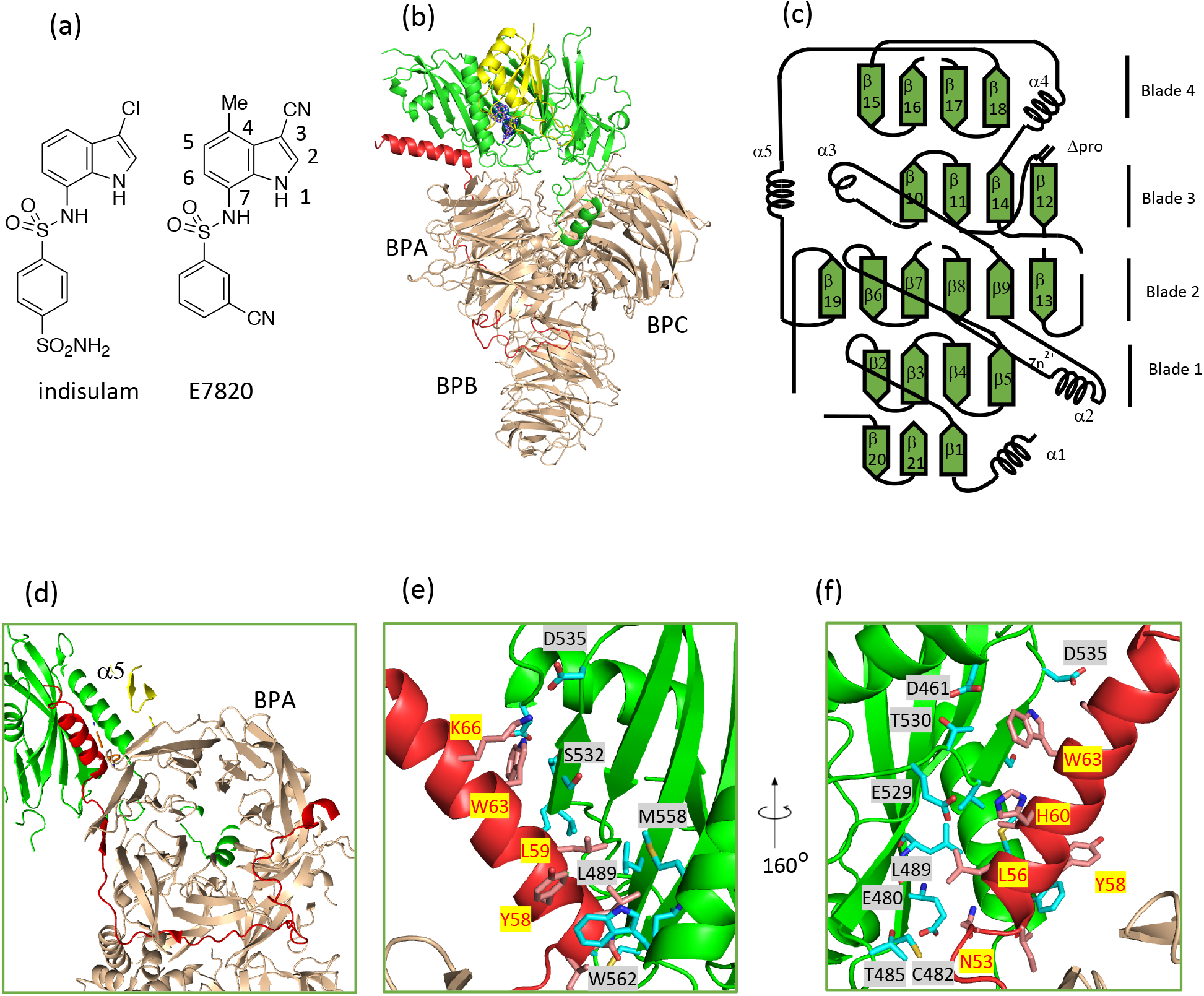
Structure of the ternary complex of DDAl-DDBl-DCAFl5Δpro-RRM2 with E7820. (a) Structures of E7820 and indisulam. (b) Overall organization of the complex of DDA1 (red), DDB1 (wheat), DCAF15Δpro (green), RRM2 (yellow), and E7820 (orange sticks and blue 2Fo-Fc electron density contoured at 1 σ). (c) Secondary structure and topology of DCAF15Δpro. (d) The organization of DDA1, BPA of DDB1, and DCAF15Δpro (looking down the barrel of BPA domain). (e) and (f) Interface between DDA1 and DCAF15Δpro at two angles.

The mechanism of action of SPLAMs is analogous to that of immunomodulatory drugs (IMiDs), such as thalidomide, lenalidomide, and pomalidomide. IMiDs bind to another adaptor protein for Cul4, cereblon, and alter its substrate selectivity to recruit, ubiquitinate, and degrade proteins including IKZF1, IKZF3, CK1α, GSPT1, and SALL4 (Ito et al. 2010; Kronke et al. 2014; Lu et al. 2014; Fischer et al. 2014; Petzold et al. 2016; Matyskiela et al. 2016; Matyskiela et al. 2018). The degrons on the neo-substrates lack a consensus primary sequence but share a common motif, an extended β-hairpin motif (Petzold et al. 2016; Matyskiela et al. 2016). RBM39, on the other hand, is expected to interact with DCAF15-SPLAM through a helix as several single mutations on the α1 helix of the RRM2 domain confer resistance to indisulam-induced degradation of RBM39 (Han et al., 2017). To better understand the structural basis of the recruitment of RBM39 to DCAF15, we determined the crystal structure of the ternary complex of DCAF15 (as a stable triplex with DDB1 and DDA1) with E7820 and RRM2. In addition, our kinetic studies on the formation of the binary and ternary complexes revealed that RBM39 binding to DCAF15 is synergistic with either E7820 or indisulam.

## Results

### Crystallization of the ternary complex

Our initial efforts to crystallize the triplex of DDA1-DDB1-DCAF15 with or without RBM39 were unfruitful. We hypothesized that the proline-rich, atrophin-homology domain of DCAF15 (amino acids 276 to380) is flexible and thereby hinders crystallization. The protein without this segment (referred to as DCAF15Δpro hereafter) is fully functional in recruiting RRM1-RRM2 domain of RBM39 (referred to as R1R2 hereafter) in an E7820-dependent manner (Supplementary Figure S1). We readily obtained crystals of DDA1-DDB1-DCAF15Δpro-R1R2 with E7820, collected a 3.0 Å data set, and solved the crystal structure. Since no electron density was observed for the RRM1 domain, we subsequently used only the RRM2 domain in the crystallization and obtained the 2.9 Å crystal structure reported here (PDB ID:6PAI). The statistics for data collection and crystallographic refinement is reported in Table 1.

**Table 1.** Statistics of data collection and structural refinement

| **Data collection** |  |
| --- | --- |
| Space group | P2<sub>1</sub> (No. 4) |
| Unit cell dimensions (Å, °) | a = 81.51, b = 94.77, c = 145.71;<br>β = 98.06 |
| Wavelength (Å) | 0.97918 |
| Average mosaicity (°) | 0.63 |
| Resolution range (Å) | 50–2.90 (2.95–2.90) |
| Unique number of reflections | 47,114 (2204) |
| Average redundancy | 3.6 (3.3) |
| Completeness (%) | 96.9 (92.3) |
| $R_{\text{r.i.m.}}$ (%)<sup>a</sup> | 10.2 |
| $R_{\text{p.i.m.}}$ (%)<sup>b</sup> | 5.2 (29.8) |
| $\langle I / \sigma_I \rangle$ | 12.3 (1.4) |
| CC<sub>1/2</sub> in the last resolution shell | 0.82 |
| Wilson $B$-factor (Å<sup>2</sup>) | 70.0 |
| **Refinement** |  |
| Resolution range (Å) | 49.5–2.90 (2.975–2.900) |
| Number of reflections (Total/$R_{\text{free}}$) | 47,085/2310 (3040/163) |
| Atoms (non-H protein) | 13,196 |
| Protein residues (resolved/sequence) | 1649/1849 |
| $R_{\text{work}}$ (%) | 20.8 (34.2) |
| $R_{\text{free}}$ (%) | 25.2 (33.9) |
| RMSD bond length (Å) | 0.003 |
| RMSD bond angle (°) | 1.13 |
| Overall $B$-factor (Å<sup>2</sup>) | 86.9 |
| Ramachandran plot (%) (favored/allowed/disallowed)<sup>c</sup> | 95.3/4.4/0.3 |
| Poor rotamers (%)<sup>c</sup> | 2.3% |
| Clashscore<sup>c</sup> | 13.6 |

Data for highest resolution shell are given in brackets.
<sup>a</sup>Redundancy-independent merging R factor,
<sup>b</sup>Precision-indicating merging R factor,
<sup>c</sup>Calculated using MolProbity (Chen et al. 2010)

### The structure of DCAF15Δpro

The overall arrangement of the complex is shown in Fig. 1b. As established in previous structural studies, DDB1 consists of three β-propellers, namely BPA, BPB, and BPC (Angers et al. 2006). BPB domain mediates the binding to CUL4 while the two rings of BPA and BPC form a half-open clamp to interact with DCAFs. DCAF15 is anchored to BPC through its N-terminal helix. A similarly positioned helix has been found in many DCAFs and viral proteins that functionally mimic DCAFs to mediate the interaction with DDB1 (Li, et al. 2010). Unlike DCAF1, another adapter protein for Cul4-related E3 ligases with known structure (Schwefel et al. 2014), DCAF15 does not fold as a seven-bladed β-propeller, like a typical WD40 protein. Instead, DCAF15 consists of five β-sheets arranged in an open, horse shoe-shaped configuration (Fig. 1b and c). The core of the fold consists of four blades that are topologically similar to the blades in a typical WD40 protein with regard to both the connections between the β-strands within a blade and the connections between the blades (Smith et al. 1999). Blade 1 and 4 have four antiparallel β-strands as in a typical blade. Blade 2 includes the topologically conserved strands β6, β7, β8, and β9 and two noncanonical strands, β 13 and β 19, which act as connecting structures. A zinc finger motif was identified between β8 and β9. The first two zinc chelating residues (C193 and C196) are positioned on the N-terminus of α2 and the second two (C211 and H214) are positioned on the subsequent loop connecting α2 and β9. Blade 3 consists of α3, β10, β 11, and β14, which are topologically conserved, and a noncanonical β 12. Interestingly, the first strand in Blade 3 morphs into a short helix (α3). The main chain NH groups of A234 and F235 at the start of the helix are free from intra-helical hydrogen bonding and thus are available to form hydrogen bond with E7820, as described later.

### Structure of DDA1

DDA1 was identified along with DDB1 and DCAF15 as part of the E3 complex that recruited RBM39 in HCT116 cells (Olma et al. 2009; Han et al. 2017). While co-expression with DDB1 was sufficient for the expression and purification of DCAF15Δpro in insect cells, further inclusion of DDA1 increased both the yield and the stability of the protein complex. Previously, N-terminal portion of DDA1 was found to bind to BPA in a crystallographic study (Shabek et al. 2018). However, only 19 residues were modeled because of the poor electron density for the rest of the chain. In this study we observed a contiguous density corresponding to amino acids 2 through 70. Much of the DDA1 wraps around BPA propeller of DDB1 while only making minimal van der Waals contacts with BPB and BPC propellers (Fig. 1d). C-terminus of DDA1 (residues 53-70) forms a long α-helix that is sandwiched between the top of BPA of DDB1 and the Blade 4 and α5 helix of DCAF15 (Fig. 1d and e). Thus, DDA1 further stabilizes the interaction between DDB1 and DCAF15.

### Interface between E7820, DCAF15Δpro, and RBM39

E7820 packs into a shallow pocket on the surface of DCAF15 (Fig. 2a), which is formed by R552, V556, V559, and M560 of α5 helix on one side and T230, Q232, and F235 on the other side (Fig. 2b). The two sulfonamide oxygens form hydrogen bonds with the main chain NH groups of A234 and F235 with good geometry. In addition, there exists a hydrogen bond between the indole NH and the main chain carbonyl oxygen of F231. The benzonitrile ring of E7820 is positioned to form a potential T-shaped π–π stacking interaction with F235 of DCAF15 (McGaughey et al. 1998; Brinlinski 2018). As expected from the results of the forward genetic study, RRM2 domain interacts with DCAF15 and E7820 mainly through its α1 helix (Han et al. 2017). We note several potential hydrogen bonds between the C-terminus of α1 helix of RRM2 and DCAF15 side chains: main chain carbonyl oxygen of G268, I269, and P272 form hydrogen bonds with side chains of R552, S549, and Y226 of DCAF15, respectively. There is good electron density to support the existence of a salt bridge between E271 of RRM2 and R178 of DCAF15 (Fig. 2c). The N-terminus of the α1 helix forms van der Waals contact with the five-membered ring of the indole group. In addition, M265 on the α1 helix appears to form interaction with the indole ring of E7820. The stabilizing role of methionine sulfur-aromatic interaction has been previously discussed (Valley et al. 2012).

**Figure 2.**
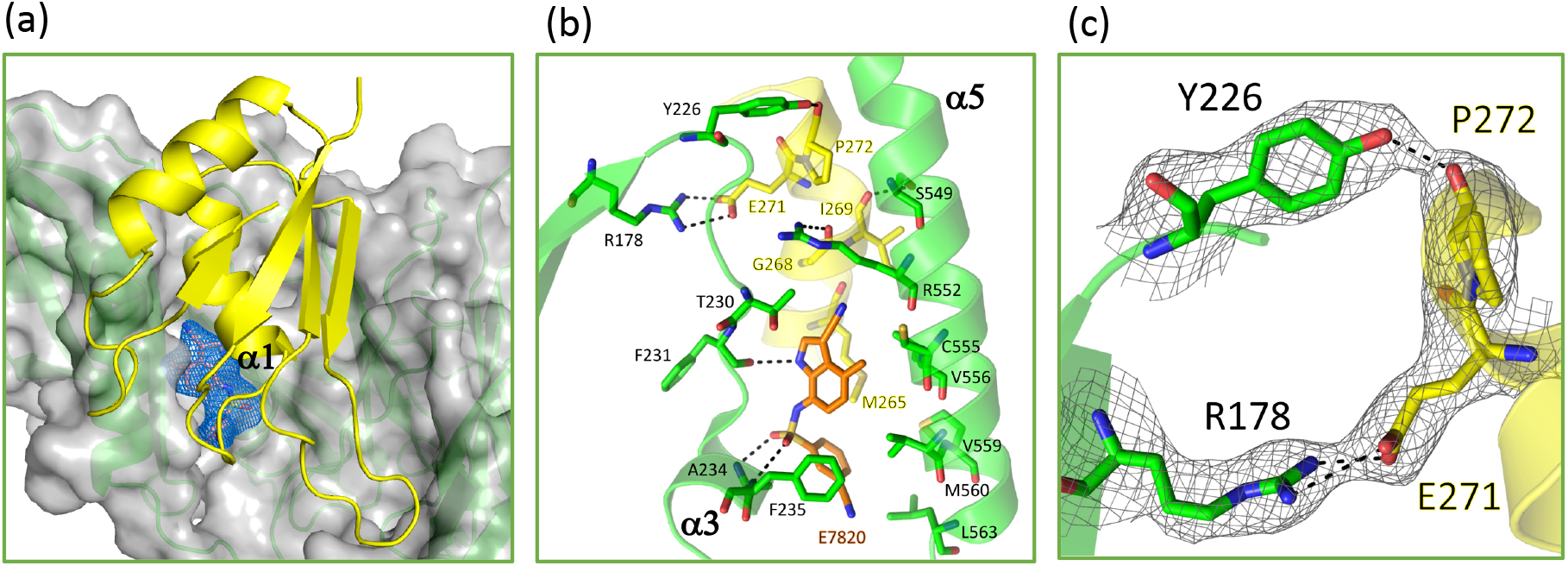
Interface between DCAF15Δpro, E7820, and RRM2. Models for DCAF15Δpro, RRM2, and E7820 are colored in green, yellow, and orange, respectively. (a) Overall view of E7820, DCAF15Δpro, and RRM2 with transparent surface of DCAF15Δpro in gray and 2Fo-Fc map of E7820 contoured at 1 σ in blue. (b) interactions in the interface in ball-and-stick model. (c) Electron density for residues involved in two specific interactions: hydrogen bonds between the carbonyl oxygen of P272 of RRM2 and hydroxyl group of Y226 of DCAF15Δpro and a salt bridge between E271 of RRM2 and R178 of DCAF15Δpro.

It has previously been shown that the following mutations on α1 helix of RRM2 abolish recruitment of RBM39 to DCAF15: M265L, G268V/W/R/E, E271Q/G, and P272S (Han et al. 2017). The structure readily provides an explanation for how these mutations block RBM39 recruitments. M265 of α1 packs against the indole ring of E7820 and provide binding energy through sulfur-aromatic interactions. Mutation to a smaller residue like leucine would decrease this interaction and destabilize the ternary complex. G268 is in tight contact with indole ring of E7820 and T230 of DCAF15, and anything larger than alanine at this position would a create steric clash with either of the binding partners. E271 forms a salt bridge with R178 of DCAF15 (Fig. 2c and d) and removal of the negative charge in E271Q mutant would impair the interaction. P272 introduces a kink into the C-terminus of α1 helix (Fig. 2c). P272S would result in a straight helix, leading to steric clash with α5 helix of DCAF15.

#### Kinetic studies

In order to understand the kinetic pathway leading to the formation of the ternary complex of DCAF15, SPLAM, and RBM39, we characterized the binary binding reactions between the three components. No interaction was detected in isothermal calorimetry (ITC) (Pierce et al. 1999) experiments when R1R2 was titrated to E7820 or indisulam (data not shown). We observed weak affinity between R1R2 and DCAF15 using both ITC and bio-layer interferometry (BLI) (Abdiche et al. 2008) and the results are shown in Fig. 3. Dissociation constant (K_d_) by these two methods agreed very well with each other (6.4 μM in ITC and 4.6 μM in BLI). Interestingly, E271Q and M265L mutations, both of which impair SPLAM-induced recruitment of RBM39, have different effects on the intrinsic affinity between R1R2 and DCAF15 (Fig. 3c and d). E271Q abolishes the binding of R1R2 to DCAF15 while M265L has a minimal effect on K_d_. This observation is consistent with the crystal structure, which reveals that E271 is directly engaged with DCAF15 by forming a salt bridge with R178 of DCAF15 while the engagement of M265 is mediated through E7820. Thus, the differential effect of the two mutations provides experimental confirmation that the interactions observed between RRM2 and DCAF15 in the crystal structure are responsible for their intrinsic affinity. To characterize binding between SPLAM and DCAF15, we synthesized PT7795, which is an analogue of indisulam conjugated with Alexa Fluor 647 (ALX647)(Fig. 4a). The binding of PT7795 to His-tagged DCAF15 can be detected by the time-resolved Förster resonance energy transfer (TR-FRET) signal between ALX647 and europium-labeled anti-His antibody. PT7795 binds to DCAF15 with a K_d_ of 0.36 μM (Fig. 4b). We developed a competitive assay using PT7795 as a probe and determined that E7820 and indisulam bind to DCAF15 with K_d_s of 22 μM and 108 μM, respectively (Fig. 4c and d).

**Figure 3.**
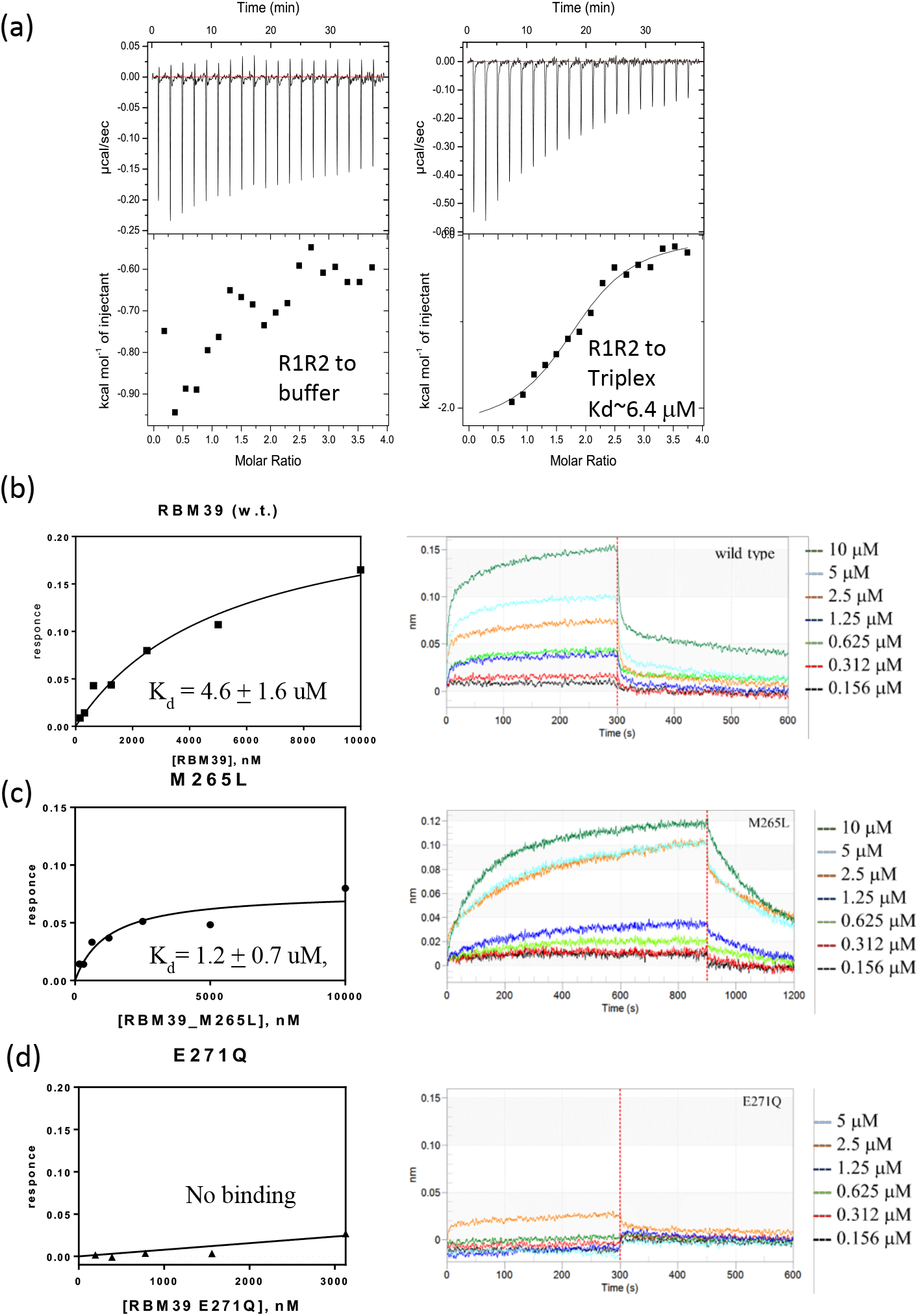
Characterization of R1R2 binding to DCAF15 by ITC and BLI. (a) Titration R1R2 to buffer or DDA1-DDB1ΔB-DCAF15Δpro triplex in ITC experiment. (b) Binding of wild type R1R2, R1R2(M265L), or R1R2(E271Q) to DDA1-DDB1ΔB-DCAF 15(Δpro) triplex in a BLI experiment. Wild type R1R2 and R1R2(M265L) bind to triplex with K_d(wt)_ = 4.6 + 1.6 uM and K_d(M265L)_ = 1.2 + 0.7 uM, respectively. No binding is detected between R1R2(E271Q) and the triplex.

**Figure 4.**
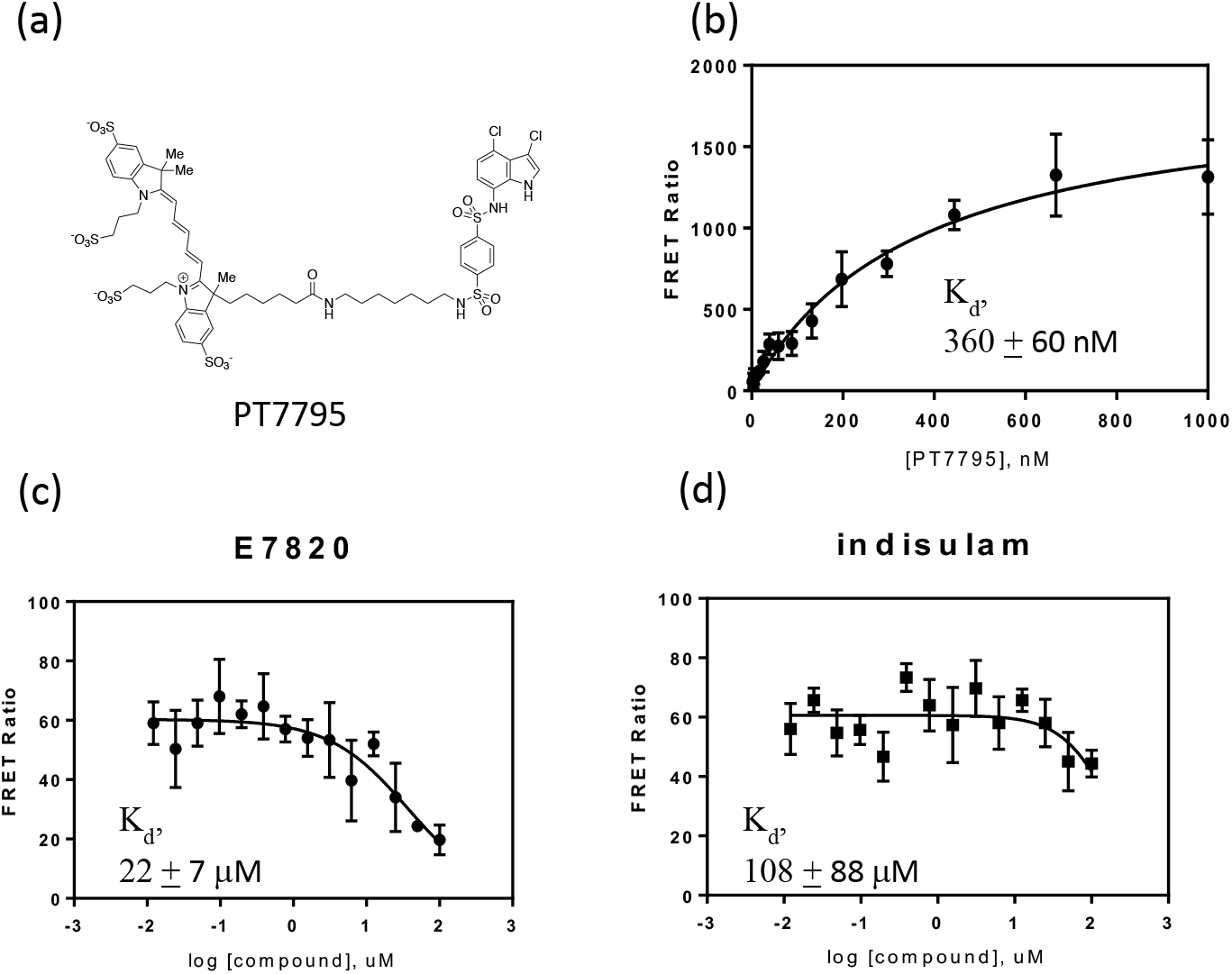
Characterization of binding of E7820 and indisulam to DCAF15. (a) Structure of PT7795. (b) FRET signal in the titration of the PT7795 to DDA1-DDB1ΔB-DCAF15Δpro. (c) and (d) Displacement of the PT7795 by E7820 and indisulam, respectively, in the competition assay. The K_d_ determined for E7820 and indisulam were found to be 22 + 7 μM and 108 + 88 μM, respectively.

Our studies on the formation of the binary complexes provide insight into the kinetics of the formation of the ternary complex. DCAF15 is able to bind a SPLAM or R1R2 independently, albeit weakly. By contrast, there is no detectable binding affinity between a SPLAM and R1R2. Therefore, the ternary complex is likely to form through two alternative pathways. DCAF15 and R1R2 can first form a weak complex, which is strengthened by the binding of a SPLAM. Alternatively, a SPLAM binds to DCAF15 first to form the DCAF15-SPLAM complex, which then recruits RBM39.

We then characterized the binding of E7820 or indisulam to DCAF15-RBM39 in a TR-FRET assay (Fig. 5a and b). His-tagged DCAF15 and FLAG-tagged R1R2 were used in the assay and SPLAM-induced complex formation was monitored through resonance transfer between europium-labeled anti-His antibody and anti-FLAG antibody conjugated to allophycocyanin (APC). As described earlier, E7820 and indisulam bind to DCAF15 weakly, with K_d_s of 22 and 108 μM, respectively. At the experimental condition with 10 nM DCAF15 and 50 nM R1R2, the apparent EC50s for ternary complex formation are 3nM and 5 nM for E7820 and indisulam, respectively. We note that these values are likely higher than the true binding constants of E7820 or indisulam to DCAF15-RBM39 because the complex was not preformed at the low concentrations of DCAF15 and R1R2 used in the experiments. Mutations of M265L and E271Q greatly reduced the formation of the ternary complex based on the magnitude of the FRET signal and the increased EC50 numbers (Fig. 5c and d). We also characterized the binding of R1R2 to DCAF15-SPLAM complex using BLI (Fig. 5e and f). The presence of a SPLAM (10μM) greatly increased the affinity of R1R2 to DCAF15 as the K_d_s decreased from about 4.6 μM to 0.16 μM and 0.12 μM in the presence of E7820 and indisulam, respectively.

**Figure 5.**
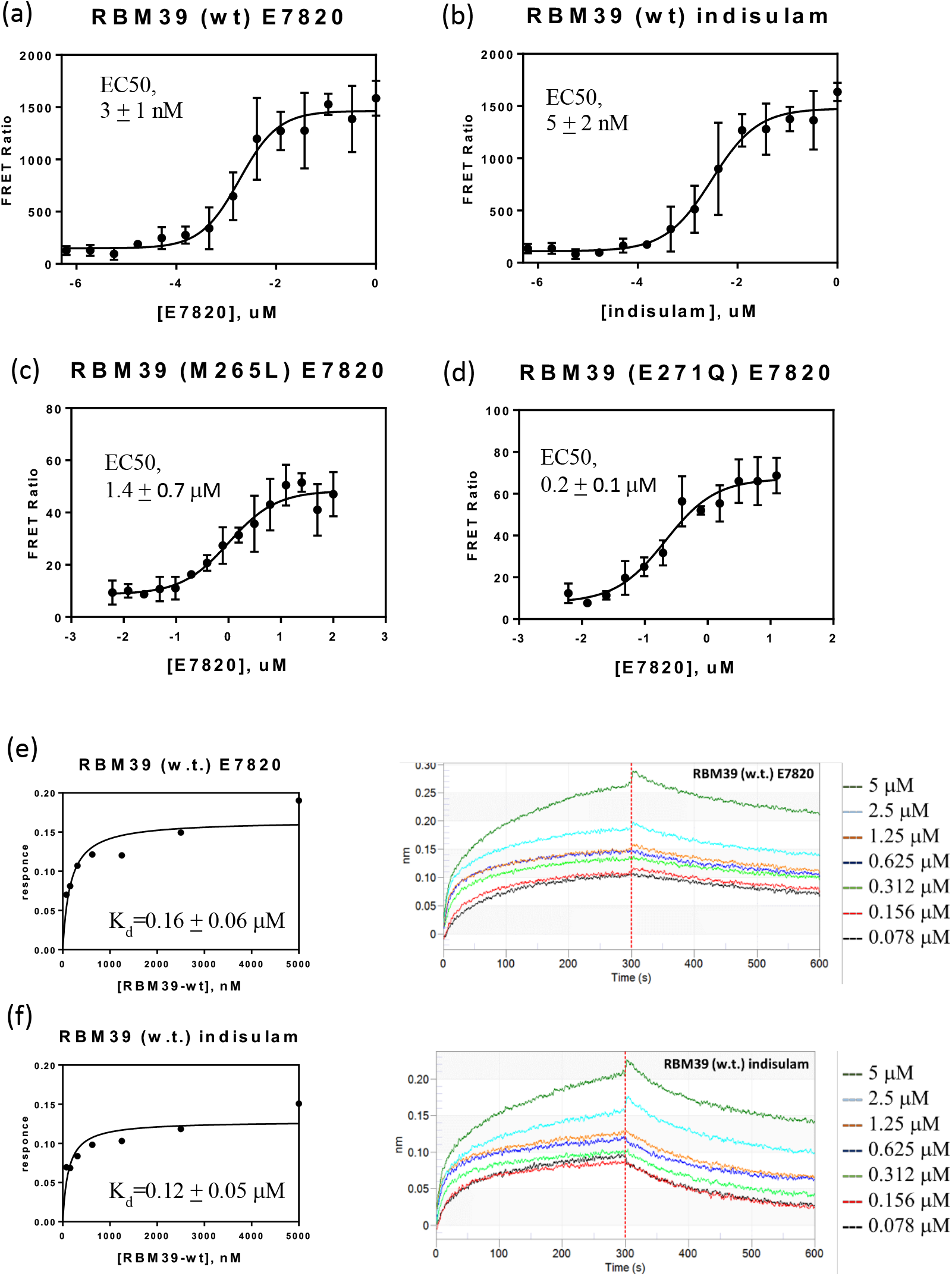
Kinetic analysis on the formation of the ternary complex. (a) and (b) FRET signal in the titration of E7820 or indisulam, respectively, to DDA1-DDB1ΔB-DCAF 15Δpro triplex (10nM) and R1R2 (50 nM). (c) and (d) FRET signal in titration of E7820 to the mixture of triplex and R1R2(M265L) and R1R2(E271Q), respectively. (e) and (f) BLI signal in the binding of R1R2 to DCAF15 in the presence of E7820 (10 μM) or indisulam (10 μM), respectively.

The observed synergy of binding of SPLAM and RBM39 to DCAF15 can be rationalized by examination of the structure of the ternary complex. The binding of the α1 helix of RRM2 to DCAF15 creates a much deeper binding site for SPLAM, enabling more van der Waals interaction and hydrophobic interaction between E7820 and the N-terminus of the α1 helix. Conversely, the same interactions strengthen RBM39 binding to DCAF15, which in the absence of a SPLAM is only mediated by the C-terminus of the α1 helix. The mechanism of the synergistic binding is schematized in Fig. 6.

**Figure 6.**
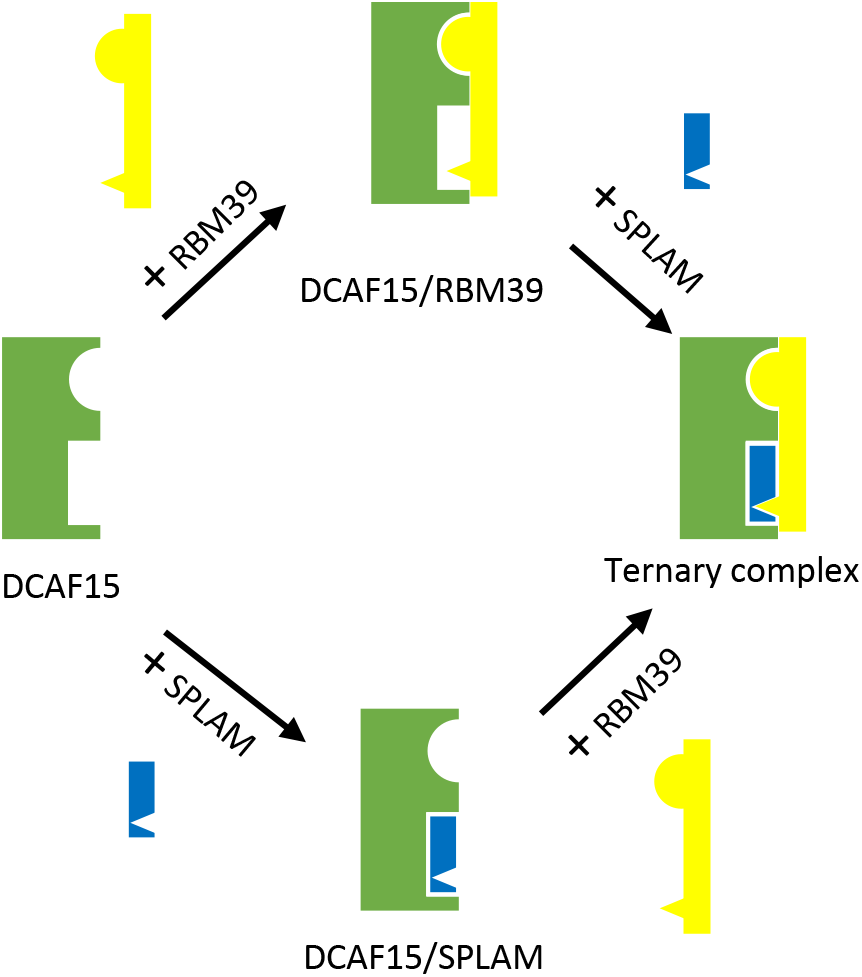
Schematics depicting two alternative pathways for the formation of the ternary complex and the structural basis of synergistic binding of RBM and SPLAM to DCAF15. Binding of either RBM39 (upper pathway) or SPLAM (lower pathway) creates additional binding surface for the other entity, thereby resulting in synergistic binding.

## Discussion

Since its identification as the target of SPLAMs, RBM39 has emerged as an intriguing cancer target. In a recent study, dependencies of 490 RNA-binding proteins were systematically interrogated in several types of cancer cells including acute myeloid leukemia (Wang et al. 2019). CRISPR-mediated deletion or pharmacological degradation of RBM39 led to preferential killing of AML cells. The presence of spliceosomal gene mutations was found to be an important predictor of response to SPLAMs in addition to the expression level of DCAF15, which was identified earlier (Han et al. 2017). Thus, while SPLAMs so far demonstrated only modest efficacy in clinical trials, patient selection based on these biomarkers might increase the potential therapeutic utility of SPLAMs. Our crystal structure of the ternary complex provides insight into how to design SPLAMs with better efficacy or drug-like properties.

In recent years small molecule-induced proteasomal degradation has emerged as an attractive strategy for drug discovery. One approach is to utilize so-called PROteolysis TArgeting Chimeras (PROTACs), which are bifunctional small molecules capable of binding to an E3 ligase with one functional group and a target protein of interest with the other functionality (Hughes et al. 2017; Pettersson et al. 2019; for reviews). Induced proximity of the target to the E3 ligase often leads to poly-ubiquitination and degradation of the target. Despite growing interest in using more E3 ligases (Ottis et al. 2017), so far only eight E3 proteins out of about 600 potential E3 ligases have been utilized in this approach, namely VHL, cereblon, cIXP, XIAP, Keap1, RNF4, RNF114, and MDM2 (Paiva et al. 2019). The primary reason is the scarcity of E3 proteins with well-characterized small ligands that can be used as a recruiting element. Our structural and kinetic studies pave the way to the development of DCAF15 and sulfonamides as a new PROTAC system. Guided by the crystal structure, we were able to synthesize PT7795, which binds to DCAF15 with sub-microMolar affinity. The fact that we were able to add large functionality (in this case, ALX647) via a linker to the phenyl ring suggests that bifunctional degraders based on sulfonamides may be achievable.

SPLAMs represent the second group of molecular glues, after IMiDs, that recruit neo-substrates to an E3 ligase. It is therefore important to note both the distinctions and common features between the two systems. As alluded earlier, IMiDs modify the surface of cereblon to recruit neo-substrates via a β-hairpin structure motif on the neo-substrates. The interface is dominated by hydrogen bonds between the main-chain carbonyl oxygens of the β-hairpin and the side chains of cereblon (Petzold et al. 2016; Matyskiela et al. 2016). Consequently, there is little requirement for sequence specificity in the β-hairpin except at a glycine position. In fact, a fairly large number of Zn-finger proteins were found to be recruited to cereblon by several IMiDs (Sievers et al. 2018). On the other hand, RBM39 is recruited through an α-helix, which interacts with DCAF15-SPLAM through both main-chain carbonyl oxygens and side chains.

The interactions are specific and several single mutations at key positions in RBM39 severely impair the recruitment. Apparently the two systems also differ kinetically in the formation of the ternary complex. In the case of cereblon, IMiDs bind first with high affinity (8.5 nM) and the binding of a neo-substrate follows (Ito et al. 2010). There was no report in the literature whether neo-substrate binding increases the affinity of an IMiD. On the other hand, DCAF15 has low and comparable affinity to both RBM39 and SPLAMs, and synergistic binding of the two leads to the formation of the ternary complex. However, we note the presence of both E3-substrate and glue-substrate interactions in the crystal structures of both systems (Petzold et al. 2016; Matyskiela et al. 2016). This common feature strongly implicates synergistic binding in both systems.

Our structural and biochemical studies provide a framework for future efforts to develop molecular glues that recruit other proteins of interest to DCAF15. As discussed earlier, the recruitment of RBM39 is highly specific. While this feature presents challenge for rational approach to the discovery of neo-substrates, it also offers opportunities to obtain highly specific molecular glue. The binding pocket for E7820 is relatively flat and is able to accommodate sulfonamides with large chemical diversity, which presents opportunity for unbiased approach to identify potential neo-substrates. Though an exciting approach to drug discovery, PROTACs have been limited to targets which already have a small molecule ligand or inhibitor. As such, many therapeutically important proteins that lack small molecule binding sites, such as most transcription factors, remain intractable by the PROTAC approach. Development of molecular glues, which rely on secondary structural features of target proteins and not on the classical ligand-binding pocket or cleft, represents a promising way to overcome this limitation and expand the druggable space in the proteome.

## Materials and Methods

### Protein Constructs and Expression

Human DDA1 (UniProt Q9BW61), DDB1 (Q16531), and DCAF15 (Q66K64) coding sequences were cloned into pFastBac1 vectors and were co-expressed in Sf9 cells using Bac-to-Bac baculovirus expression system (Thermo Fisher Scientific). The expression construct for DCAF15 includes a N-terminal His6-tag to facilitate the purification. To improve the crystal quality, the atrophin-1 homology region of DCAF15 (amino acid residues 276 – 383) was excised. While the triplex of DDA1-DDB1-DCAF15Δpro was useful for obtaining diffracting crystals, triplex of DDB1ΔB, which has the BPB domain (aa 396 to 705) excised and replaced with a GNGNSG linker, was used in all the kinetic measurements because of the higher yield. R1R2 of RBM39 (aa 150 to 331) and the mutants thereof were expressed as GST-fusion proteins. The coding sequence was sub-cloned into pGEX4T3 with an engineered N-terminal TEV protease cleavage site. A C-terminal FLAG tag was also added to R1R2 used in FRET assay. RRM2 domain of RBM39 (amino acids 235-331) was sub-cloned directly into pET28a and expressed with a N-terminal His-tag.

### Protein Purification Summary

Detailed information of protein purification can be found Supplementary Information. In short, the triplex of DDA1-DDB1-DCAF15ΔPro was purified through sequential application of Ni affinity (Profinity, Bio-Rad), size-exclusion (HiLoad 16/600 Superdex 200 pg, GE Healthcare), and ion exchange (HiTrap Q, GE Healthcare) column chromatography. GST-fused R1R2 or mutant was expressed in *E coli* and purified using agarose-glutathione beads. R1R2 was released from GST with TEV protease treatment and furthered purified by ion exchange chromatography using a HiTrap SP column. His-tagged RRM2 domain of RBM39 was expressed in *E coli* and purified using Ni^2^+-charged affinity resin. His-tag was then removed with thrombin and RRM2 was further purified by ion exchange chromatography using a HiTrap Q column. To obtain the tetraplex of DDA1-DDB1-DCAF15Δpro-E7820-R1R2 (or RRM2), R1R2 (or RRM2) was added to the triplex with two-fold Molar excess and E7820 was added to 100 μM. The mixture was then loaded onto tandem size-exclusion columns (Superdex 75 and Superdex 200, 10/300), that were equilibrated with 20 mM Tris-HCl, pH8.0, 150 mM NaCl, 1mM TCEP, and 10 μM E7820. The peak containing DDA1-DDB1-DCAF15-E7820-R1R2 (or RRM2) was concentrated to 4.0 mg/mL for crystallization trials.

### Crystallization and Structural Determination

Crystallization screens for the complexes were set up on a Gryphon robot (Art Robbins) and incubated at 4 °C. Single crystals for DDA1-DDB1-DCAF15Δpro-E7820-R1R2 were obtained with condition F2 of Proplex screen (0.2 M lithium chloride, 0.1 M Tris-HCl, pH 8.0, and 20% PEG 8K, Molecular Dimensions). The freezing solution consisted of the same buffer components and 30% glycerol. A 3.0Å data set was collected at Beamline 821 at ALS. Single crystals for DDA1-DDB1-DCAF15Δpro-E7820-RRM2 were obtained with condition B11 of Proplex screen (0.2 M HEPES, pH7.5, and 20% PEG 4K). A 2.9 Å data set was collected at beamline I.D. 19 at the Advanced Photon Source, Argonne National Laboratory. The structure of DDA1-DDB1-DCAF15Δpro-E7820-RRM2 was solved by molecular replacement using Phaser (McCoy et al. 2007), with BPA-BPC (4AOL), BPB (4AOL), and RRM2 (2JRS) as searching models. DCAF15 and DDA1 models were manually built in Coot (Emsley et al. 2004) alternating with positional and isotropic atomic displacement parameter (ADP) refinement in REFMAC5 (Vagin et al. 2004; Murshudov et al. 2011). Both isotropic ADP and TLS group refinement were used (Winn et al. 2001; Painter et al. 2006). The geometry restraints for E7820 were generated in AceDRG (Long et al. 2017). Refined structures were analyzed in MolProbity (Chen et al. 2010). Atomic representations were created using MacPyMOL (Version 1.5.0.3, Schrödinger).

### ITC

ITC experiments were carried out on an iTC200 (Malvern). The triplex of DDA1-DDB1ΔB-DCAF15Δpro and R1R2 were first concentrated to about 0.04 mM and 0.75 mM, respectively. The two samples were dialyzed against the buffer consisting of 20 mM Tris-HCl, pH 7.6, 150 mM NaCl, and 1 mM TCEP at 4 °C overnight. R1R2 at 0.75 mM was titrated into the triplex at 0.04 mM at 20 °C and the thermogram was fitted using a one binding site model.

### BLI

BLI experiments were carried out on an Octet RED96e system (ForteBio). Octet Buffer consisted of 25 mM HEPES pH7.5, 100 mM NaCl, 0.1mg/ml BSA, and 0.005 *%* Tween 20. In the loading step, Ni (NTA) biosensors were dipped into wells containing His-tagged DCAF15Δpro-DDA1-DDB1ΔB at 12.5 μg/mL for 300 seconds. Equilibration of biosensor was achieved by dipping sensors into buffer alone for 120 seconds. The association step involved dipping the probes into wells containing either R1R2 alone or R1R2 with SPLAM at 10 μM for 300 to 900 seconds. The dissociation step was measured by next dipping the biosensors into wells containing buffer alone and or buffer with SPLAM at 10 μM for 300 seconds. Resulting data was either fit globally or locally to a one step binding model. K_d_s were determined by fitting the steady state response to equation 1.

### Binding of E7820 and indisulam to DCAF15

Binding of PT7795 to DCAF15 was measured by the FRET signal between europium-labeled anti-His antibody (AD0205, Perkin Elmer) that is associated with His-tagged DCAF15 and ALX647 moiety in PT7795. The concentration of PT7795 was varied from 2.3 nM to 1000 nM in 16 wells, each containing 25 mM HEPES pH7.5, 100 mM NaCl, 0.1mg/ml BSA, 0.005 % Tween 20, 0.5 mM TCEP, 4% DMSO, 0.5 nM europium-labeled anti-His antibody, and 10 nM triplex of DDA1-DDB1ΔB-DCAF15Δpro. After equilibrating for 2-3 hours at 4 °C, the plate was removed from the incubator, briefly spun, then read on a Spark 10 (Tecan) at the excitation/emission wavelength of 340nm/615nm and 340nm/665nm. To obtain K_d_, the ratio of 665/615 readings was fitted to Equation 1, where *F* stands for the ratio of 665/615 nM.

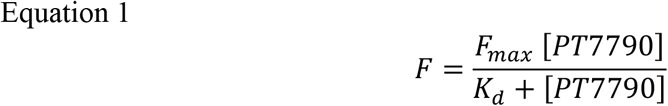

Binding of E7820 and indisulam to DCAF15 was measured in a competition by the displacement of PT7795. The concentration of PT7795 was kept at 300 nM while the concentration of E7820 or indisulam was varied from 0.01 to 100 μM. The incubation time was increased to 18 hours for enhanced signal to noise level. To obtain IC_50_, the ratio of 665/615 readings was fitted to Equation 2, where n is Hill slope.

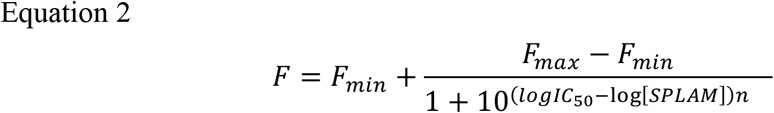

The K_d_s were then calculated from IC_50_s using Cheng-Prusoff equation (Cheng et al., 1973):

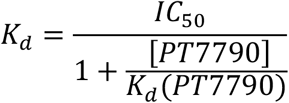

### DCAF15-RBM39 complex formation monitored by time-resolved FRET analysis

His-tagged DDA1-DDB1ΔB-DCAF15Δpro triplex and R1R2 with a C-terminal FLAG were used in the assay and SPLAM-induced complex formation was monitored through resonance transfer between europium-labeled anti-His antibody and APC-labeled anti-FLAG antibody (AD0059F, Perkin Elmer). Assay buffer contained 25 mM HEPES, pH7.5, 100 mM NaCl, 0.1mg/ml BSA, 0.005 % Tween 20, 0.5 mM TCEP, 0.5 nM europium-labeled anti-His antibody, 50 nM APC-labeled anti-FLAG antibody, 10 nM triplex of DCAF15, and 50 nM R1R2. The concentration of the SPLAM was varied and the ratio of 665/615 nm readings was fitted to Equation 2 to obtain the apparent of EC50 for SPLAM-induced complex formation.

## Acknowledgements

This research used resources of the Advanced Light Source, which is a Department of Energy (DOE) Office of Science User Facility under contract no. DE-AC02-05CH11231 and Advanced Photon Source, a DOE Office of Science User Facility operated by Argonne National Laboratory under Contract No. DE-AC02-06CH11357.

## Author information

### 3. Contributions

X.D. designed and made constructs for recombinant protein expression. O.V. and H.T. prepared the viruses for DDA1, DDB1, DDB1ΔB, DCAF15, DCAF15Δpro. H.T. worked out the procedure for the purification of DDB1ΔB-DCAF15Δpro complex. O.V. purified the DDA1-DDB1-DCAF15ΔPRO complex for crystallization and the DDA1-DDB1ΔB-DCAF15ΔPRO for biophysical assays. X.D. purified R1R2 and RRM2, prepared the tetraplexes, and set up screens for crystallization. X.D., C.H., and O.V. were involved in optimization of the crystals. X.D. collected the data and processed the data. X.D. and O.V. solved the structure and built the models. O.V. refined the structure. R.S. and E.M. designed and carried out the FRET and BLI experiments. X.D. carried out the ITC experiments. P.W. and R.X. synthesized PT7795. X.D. wrote the manuscript. X.D., J.R., and E.W. oversaw the project.

## Supplementary information

### Protein Purification

Cell pellets were collected 65 h post-infection by centrifugation at 4000 × g for 5 min and suspended in cold lysis buffer (20 mM Tris-HCl, pH 7.6, 200 mM NaCl, 10% (v/v) Glycerol, 1% (v/v) Nonidet P40 substitute, 1 mM tris(2-carboxyethyl)phosphine (TCEP)) supplemented with EDTA-free Pierce protease inhibitor tablets (Thermo Fisher Scientific), 35 mL per liter of culture. All further steps were performed at 4 °C or on ice. The cell suspension was lysed by sonication (2 s on, 2 s off, for 10 min on ice) and cleared by centrifugation at 35,000 rpm, 1 h. The supernatant was incubated with Ni^2+^-charged affinity resin (Profinity, Bio-Rad) for 1 h, 1.5 mL of resin per liter of culture. The resin was washed with 10 column volumes (CV) of Wash buffer (20 mM Tris-HCl, pH 7.6 (at room temperature), 200 mM NaCl, 5 mM imidazole, 1 mM TCEP), and protein was eluted with 5 CV Elution buffer (Wash buffer supplemented with 250 mM imidazole). The eluted protein sample was concentrated by ultrafiltration (30,000 NMWL, Amicon, Millipore Sigma) and further separated on size exclusion chromatography column (HiLoad 16/600 Superdex 200 pg, GE Healthcare) connected to Akta FPLC (GE Healthcare Life Sciences) using GF buffer (20 mM Tris-HCl, pH 7.6 (at room temperature), 150 mM NaCl, 1 mM TCEP) as mobile phase. The presence and stoichiometry of the three proteins in a UV absorbance peak approximately corresponding to the expected size of the complex (195 kDa for full-length DDB1-containing complex and 160 kDa for DDB1ΔB-containing complex) was confirmed by SDS-PAGE. The protein-containing fractions were concentrated as above and diluted with QA buffer (20 mM Tris-HCl, pH 7.6 (at room temperature), 1 mM TCEP) to 30-40 mM NaCl final. The protein sample was then separated by ion-exchange chromatography using 1 mL HiTrap Q HP column (GE Healthcare Life Sciences) and a gradient of 0-1 M NaCl in QA buffer over 50 mL. The complex eluted with 400 mM NaCl and was concentrated as described above and diluted with QA buffer to 50-100 mM NaCl. The samples were flash-frozen and stored at 5-10 mg/mL protein concentration at −80 °C.

**Figure S1.**
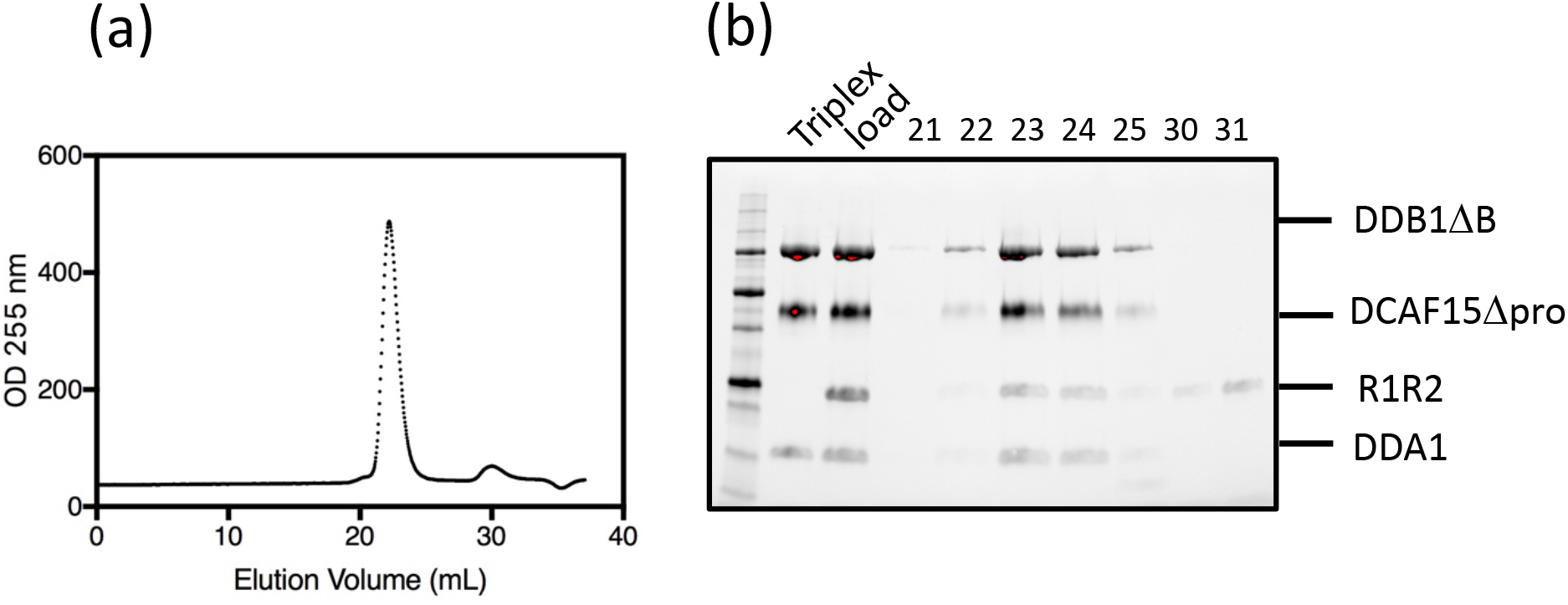
Triplex of DDAl-DDBl△B-DCAFl5△pro co-elutes with R1R2 on size-exclusion column. Chromatogram and SDS-PAGE gel of the fractions are shown in (a) and (b), respectively. “Triplex” lane contained the complex of purified DDAl-DDBl△B-DCAFl5△pro. The “load” lane contained triplex with R1R2 in two-fold Molar excess. The buffer for the size-exclusion column chromatography consisted of 20 mM Tris, pH8.0, 200 mM NaCl, 1mM TCEP, and 10 μM E7820.

